# Correlated evolution of the neck, head and forelimb across the theropod-bird transition

**DOI:** 10.1101/2025.04.04.647043

**Authors:** Ryan D. Marek, Ryan N. Felice

**Affiliations:** Centre for Integrative Anatomy, Department of Cell and Developmental Biology, University College London, London, UK; Natural History Museum, London, UK; Department of Genetics, Evolution and Environment, University College London, London, UK

## Abstract

Powered flight has required birds to undergo numerous dramatic and coordinated evolutionary responses across the entire body, yet studies are limited to a small number of traits and often exclude a critical component of the vertebrate skeleton – the vertebrae. The neck is a critical region of the avian spine as it operates in tandem with the head as a surrogate forelimb across a diverse array of behaviours. However, the drivers of cervical vertebral evolution remain poorly understood. Here, we model shifts in adaptive optima and evolutionary rates of the neck, forelimb and head of extinct dinosaurs and extant birds to test if these modules co-evolved. We observe a co-occurrence of adaptive optima shifts for neck and forelimb proportion at the base of Avialae – to vertebrae adapted for stability and a forelimb better adapted for flight. These patterns are due to shifts in neck and forelimb allometry and suggest that heterochrony is an important factor in avian neck and forelimb evolution. Further, we find lower rates of both neck and forelimb evolution in birds compared to their non-avian theropod ancestors. The coordinated evolutionary response of the neck and forelimb is a derived feature of Avialae that initially evolved to stabilise in-flight vision. This axio-appendicular co-evolution has contributed to avian macroevolutionary dynamics by facilitating the evolution of a novel locomotory mode without sacrificing the grasping capability needed to directly interact with a huge diversity of environments.

## Introduction

Powered flight is a key behavioural innovation, one that has allowed multiple clades across the tree of life to thrive in entirely new niches (1–4). Birds are the exemplars of this innovation as flight has allowed them to become the most speciose clade of extant Tetrapoda (1). Such ecological diversification is thanks to an extraordinary transformation from their iconic non-avian dinosaur antecedents that involved, among other morphological and neurosensory transformations, sustained miniaturization (5), a decoupling of fore- and hindlimb modules (6–8) and numerous changes to skull morphology and integration patterns (9–12). Research into the evolution of avian powered flight is focused on cranial (10,11) and appendicular adaptations (7,8,13), with little focus on other functional and anatomical modules, nor on the coordinated response to flight between these modules (although studies that consider multiple anatomical modules are starting to appear, see references 15–18). A key component of the skeleton that is rarely discussed in this context is the neck. In extant avians the neck is an important element of the axial skeleton that positions the head to partake in a wide variety of behaviours – from preening to tool use to even tripedal locomotion (18–23).

Over the course of theropod evolution the neck appears to have undergone a morphological shift, from a robust musculoskeletal system primarily adapted for supporting a large head and assisting in hypercarnivory (24–26), to an elongated and gracile configuration that supports a myriad of key behaviours (27–29). This transformation is theorized to be correlated with shifts in other major components of the prethoracic skeleton such as decreases in head size and the loss of a grasping-capable forelimb as powered flight evolved (16,29–32). However, studies into early avian evolution often consider morphological systems in isolation and exclude the axial skeleton entirely (7,8,10,11,13). As such, the timings and mechanisms that govern the evolution of this critical portion of the avian skeleton remain unknown. One such mechanism that is responsible the evolution of morphological adaptations is heterochrony, which is defined as the change in timing and rates of developmental processes (33,34). Heterochronic shifts to a shorter development period result in paedomorphosis, whereby the descendent clade evolves to resemble the juvenile form of the ancestral clade. Paedomorphosis is an important mechanism as it is credited as the mechanism behind many key avian innovations such as a small body size (5,35,36) and a relatively enlarged orbit and neurocranium (37). However, the contribution of heterochrony to the evolution of avian postcranial morphology, especially axial morphology, is understudied.

The potential correlated evolution of the neck, head and forelimb across the dinosaur-bird transition is theorized to result in the neck functioning as a ‘surrogate forelimb’ in extant birds, allowing the functional burden of environmental manipulation to be shifted to the neck from the forelimb (29–31). Evidence for the correlated evolution of head, neck and forelimb is present in extant avians as significant morphological integration exists between all three of these modules in many groups of birds (32). This pattern is exemplified in arboreal and aquatic birds, as integration promotes shifts in the rate of neck morphological evolution (32). Thus, the integration of the neck, head and forelimb is a keystone innovation of birds as it has facilitated a coordinated response of the prethoracic skeleton when entering new niches, allowing birds to successfully interact with new ecosystems (29,30,32). Understanding how and when this correlation between the head, neck and forelimb evolved in avians and their dinosaurian ancestors is thus a critical step in further understanding how birds came to succeed in modern ecosystems.

There are numerous examples of separate anatomical modules co-evolving during the evolution of birds (6–8,14,15,17,38) in a mosaic fashion (whereby separate anatomical modules display distinct evolutionary modes and timings (38)) and these are key features of avian macroevolution (6,8,10,14). Across the theropod-bird transition, the pelvic and pectoral locomotor modules decoupled and this promoted the co-evolution of each locomotor module with other anatomical modules (6–8). Coordinated evolution of the forelimb and tail gave rise to the flight control system exhibited by extant avians, and by emphasizing different combinations of the wing, hindlimb and tail, the diversity of avian locomotor diversity increased (6,7). Recent work has shown that in extant avians this mosaic pattern of modularity also includes the head and neck, as both affiliate with the forelimb to allow birds to succeed in arboreal and aquatic environments (32). This paper aims to build on this recent work and uncover how the potential head-neck-forelimb integration evolved in birds, and when this feature arose in avian evolutionary history. By understanding this we will gain a more complete insight into how integration and modularity promote the evolution of key innovations across the broader avian bauplan in response to the evolution of powered flight.

The correlated evolution between the head and the neck may have arisen in birds due to the reduction in head mass along the avian stem lineage (31,32). Head mass is a known constraint on neck morphology across many terrestrial vertebrates as the weight of the head must be supported by the neck at all times (39–42). As relative head size is thought to decrease as theropod diet shifts away from hypercarnivory along the avian stem lineage (43,44), it is predicted that neck morphology would be released from the constraint of head size (29,31). We hypothesise that this reduction in head size could be the cause of this head-neck integration (32). Neck-forelimb integration may have arisen alongside the loss of the forelimb’s grasping capability as powered flight evolved in early avialans (29–32). It has been proposed the neck of birds evolved due to a morphofunctional compensation that was driven by heterochrony between the loss of hand functions of theropods as the forelimb was integrated into the avian wing and as the kinematic functions of the avian beak were enhanced (30). This ‘surrogate forelimb’ hypothesis is supported our recent work on extant avians (31,32), however has yet to be formally tested by assessing the tempo and directionality of neck and forelimb evolution across the theropod-bird transition

Here we investigate how and when the unique avian neck evolved by exploring how the interplay between the evolution of the neck and other regions of the body influenced the macroevolutionary dynamics of birds and their theropod ancestors, as well as exploring the heterochronic patterns of the neck, head and forelimb. We use Bayesian Ornstein Uhlenbeck (OU) modelling approaches (45–47) to model shifts in adaptive optima as well as shifts in evolutionary rates of head, neck and forelimb allometry across 222 species of avian and non-avian theropods. We predict that concurrent shifts in the adaptive regimes of neck morphology and forelimb morphology, to shorter centrum lengths and a more distally-dominated forelimb respectively, would support the surrogate forelimb hypothesis.

Conversely, if neck evolution is primarily driven by a relaxing of head mass constraint as head size decreases along the avian stem lineage we predict that a shift to a lower head mass around the base of Maniraptora would be immediately followed by a shift in adaptive regimes of neck morphology, as well as an increase in rates of neck morphological evolution. We predict that both of these hypothesis will involve paedomorphic shifts in neck, forelimb and head allometry. Since decreases body size are an important facet of early avialan evolution due to the extensive miniaturization along the avian stem lineage we explore the degree to which the observed patterns in neck, head and forelimb morphology were related to allometric effects. We use this methodology to specifically answer whether the evolution of the avian cervical spine was correlated with the loss of a grasping-capable forelimb and if neck evolution was constrained by large head size in non-avian theropods. By investigating the correlated evolution of multiple anatomical modules this study aims to unearth how a key innovation, powered flight, led to an evolutionary cascade of adaptations across the prethoracic skeleton that negated the negative consequences of evolving a grasping-incapable wing.

## Methods

### Specimen information & data collection

We collected length measurements for the cervical centra, forelimb bones (humerus, radius, ulna, longest metacarpal element) and skull of 222 species (224 specimens) of avian and non-avian theropods (Table S1). 112 of these species were extant avians and the remaining 110 were extinct avialan and non-avialan theropods (Table S1). We measured 20 of the 110 species of theropods directly from surface scans collected with an Artec Leo 3D surface scanner from multiple North American museums (Table S1). We then measured the remaining 90 specimens from the literature (Table S1). Finally, we took avian skeletal measurements from the dataset of Marek & Felice 2023. We assessed centrum length along the ventral midline of the centrum across all specimens (27), and then calculated centrum length for extant avians using the interlandmark distance between appropriate centrum landmarks used in Marek & Felice 2023 (see Fig S1 of ref. 33). We chose length of forelimb elements and relative forelimb proportion (brachial index) as they capture the major axes of forelimb morphological variation (17) and correlate with flight capability in extant avians (48– 51). Centrum length was chosen as it correlates with measures of intervertebral flexibility (52,53) (although see ref. 28)).

### Phylogenetic Hypotheses

We assembled an informal supertree using a recent hypothesis of the phylogenetic relationship of birds based on 63k intergenic coding regions (54) as a backbone. We then grafted a time-calibrated supertree of non-neornithine birds (55) on to this backbone using the ‘tree.merger’ function in the ‘RRphylo’ R package (56). This crown and stem bird supertree was extended by grafting on a second supertree that spanned the entire Mesozoic theropod phylogeny (including early diverging avialans) (57). We added theropod and early avialan taxa that were present in our dataset but missing from the supertree at this stage by grafting individual taxa from a composite tree of Dinosauria that was pruned to include just theropods (35). We time calibrated the trees from Wang & Zhou (57) and Benson et al. (35) prior to grafting the backbone. This was achieved by using the tip dates bracketed by the first and last appearance dates of the geological stages in which each taxon was collected. We calibrated the respective trees using these first and last appearance dates using the ‘minimum branch length (‘mbl’, one million years) methods within the ‘timePaleoPhy’ function of the ‘paleotree’ R package (58). These time calibration methods were similar to those used in the original publications (57). We adapted procedures from Cooney et al. (59) that combined the backbone of major avian relationships from (54) with the fine-scale species relationships from a maximum clade credibility tree generated from www.birdtree.org (60) to produce a phylogeny that contained all the extant avian species. This final tree of 10,199 species was pruned to match the 222 species in our dataset.

### Evolutionary shifts in the adaptive optima of neck, forelimb and head morphology

We estimated adaptive shifts in cervical, forelimb and head morphology using a Bayesian multi-regime Ornstein Uhlenbeck (OU) modelling approach in the R package ‘bayou’ (45). A phylogenetic generalized least squares (PGLS) regression indicated that the lengths of the cervical centra, forelimb and head and forelimb proportion metrics all displayed a significant relationship (p < 0.001) with body mass. We therefore created custom bayou models that included body mass a predictor variable to test for shifts in the allometric scaling of cervical, pectoral and cranial morphology. We used these custom models to test specific biological hypotheses. First, we tested whether adaptive shifts to a more distally-dominated forelimb were concurrent with shifts in neck morphology at the base of Avialae, this would allow us to understand whether the evolution of the avian neck was correlated with the loss of a grasping-capable forelimb. Second, we tested whether adaptive shifts to a shorter relative head length were concurrent with shifts in neck morphology to understand if neck evolution was constrained at all by large head size in non-avian theropods. We used the ‘shiftSummaries’ and ‘plotShiftSummaries’ bayou functions to determine if adaptive shifts were heterochronic by observing if a shift in y-intercept (trait morphology) occurred between the ancestral and descendent regimes that were identified by the above bayou models.

We selected priors for the constraint parameter (α), evolutionary rate (σ^2^), and phenotypic optimum (θ) to ensure acceptance ratios remained between 0.2 and 0.4 for each parameter. We set the prior for the body mass regression coefficient (β_BM_) to 0.7, drawing on previous studies of avian neck allometry (31). Once the priors were set, we ran five independent MCMC chains and assessed convergence for each chain using the Gelman and Rubin’s R statistic (61,62). We used the combined chains resulting from this procedure to assess allometric shifts in adaptive optima of each measurement of cervical, forelimb and head morphology. For each individual dataset (first/mid/last centrum length, humerus/radius/ulna/MC2 length and head length) five MCMC chains of 1,000,000 generations were run independently and then combined, with the first 30% of each chain were removed as burn-in. We performed model selection by examining marginal likelihoods and Bayes Factors of each model (Tables S2, S3). In each case (aside for MC2), the highest likelihood model allowed for separate intercepts and global slopes where shifts have an equal probability across branches and can only occur once per branch.

### Evolutionary shifts in the rate of trait evolution of the neck, forelimb and head

We used fossil BAMM v2.5.0 (46,47) to model the variation and potential shifts in the rate of morphological evolution of neck, forelimb and head morphology. We modelled rates using a MCMC chain with 90,000,000 generations for each trait analysed. We set model priors automatically using the ‘setBAMMpriors’ function in the BAMMtools R package. We sampled parameters every 10000 iterations, the first 10% of which were deleted as burn-in. We analysed the effective sample size of the log-likelihood and number of shift events of each chains in order to assess convergence and found that all chains in this study successfully converged (effective sample size, ESS, > 200). We accounted for changes in body mass by running a second round of MCMC chains that used the residuals from phylogenetic GLS regressions of each morphological trait versus body mass. We carried out these regressions using the ‘gls’ function of the R package ‘nlme’.

### Assessing the influence of body mass on shifts in neck, head and forelimb morphology

We calculated the phylogenetic means for head, neck and forelimb morphology, as well as body mass for each of the regimes (‘descendent’ grade) found in the bayou analysis and compared them to the mean of the ancestral grade following procedures from Smaers et al. (63–65) We used this procedure to assess the extent to which body mass influenced the recovered patterns of neck, head and forelimb morphology across the shifts recovered in the bayou analysis. The ratio of the difference in ancestral-to-descendent mean trait (neck/head/forelimb) morphology to the difference in ancestral-to-descendant mean body mass is an indication of the proportionality of ancestral-to-descendant change in mean trait morphology to mean body mass. We set the expected proportionality of this ratio as the scaling coefficient of the trait-to-body relationship of the ancestral grade. We then used the upper bound of the 95% confidence interval of the ancestral grade scaling coefficient to infer that the change in mean trait morphology from ancestral-to-descendant grade is characterized by more or less change in the trait than mean body mass. This process was inversed to evaluate the mass-to-trait scaling relationship (63,64).

## Results

### Evolutionary shifts in adaptive allometric optima of neck, forelimb and head morphology

There is strong support for a singular regime shift in the adaptive allometric optima of vertebral morphology on the branch leading to Avialae in all three cervical vertebral regions studied (posterior probability = 0.70-0.82, Figs 1, S1). Across all three studied vertebral regions, avialans and non-avian theropod dinosaurs occupied distinct adaptive optima, with avialans displaying shorter cervical vertebrae relative to non-avian theropods (Figs 1, S1). The aforementioned shift in vertebral morphology is associated with a downshift in the allometric scaling of centrum length (Figs 1, S1) meaning that centrum length is lower than expected for a given body mass in avialans when compared to non-avialan theropods. The slopes of these scaling relationship does not change across any of the observed regime shifts.

**Fig 1:**
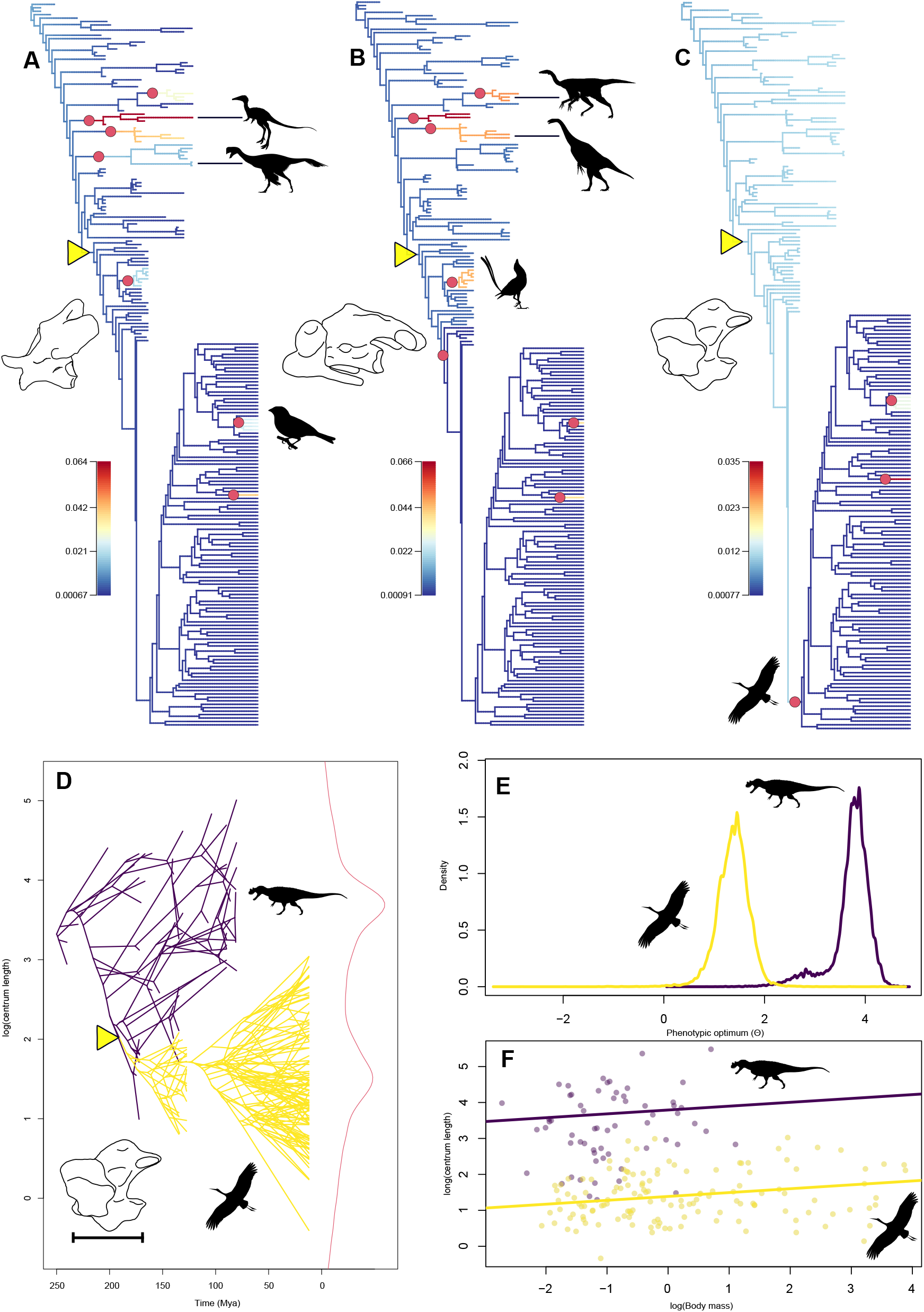
Shifts in adaptive optima (yellow triangles) and evolutionary rates (red circles) of neck morphology across non-avian theropods and avialans (including extant birds). A-C) Phylogenetic trees of non-avian theropods and avialans depicting the evolutionary rates of neck vertebral morphology (warmer branch colours denote faster rates of morphological evolution) for the first (A), middle (B) and last (C) cervical vertebrae. D) Traitgrams depicting the mode of phenotypic optimum evolution for centrum length (the measurement of which is demonstrated in the bottom left of the traitgram) evolution of the last cervical vertebrae. E) Density chart for the phenotypic optima of centrum length for non-avialan theropods (purple) and Avialae (yellow). F) Allometric scaling relationships of centrum length for non-avialan theropods (purple) and Avialae (yellow). Silhouettes are taken from PhyloPic.org and correspond to representative members of Alvarezsauroidea (*Shuvuuia deserti*), Oviraptorosauria (*Oviraptor philoceratops*, credit: Ivan Iofrida), Emberizoidea (*Emberiza sp*.), Ornithomimidae (*Struthiomimus altus*), Therizinosauria (*Nothronychus mckinleyi*, credit: Scott Hartman), Bohaiornithidae (*Bohaiornis guoi)*, Aves (*Ciconia ciconia*, credit: L. Shyamal) and Neotheropoda (*Ceratosaurus nasicornis*, credit: Scott Hartman).

Our models of humerus, radius, ulna and MC2 length scaling recovered an adaptive optimum regime shift in the branch leading to Apodiformes (posterior probability = 0.44-0.95, Fig 2). In each case, these shifts are to shorter bone lengths (Fig S2). This represents a shift to a distinct forelimb phenotypic trait optima in Apodiformes (hummingbirds and allies). The low number of Apodiformes present in this study makes this shift difficult to visualise on a phenogram (Fig 2), however the shift is well supported with high values of posterior probability (PP_humerus_ = 0.96, PP_radius_ = 0.67, PP_ulna_ = 0.70, PP_MC2_ = 0.45) and low standard error values (SE_humerus_ = 0.002, SE_radius_ = 0.003, SE_ulna_ = 0.003, SE_MC2_ = 0.002) across all forelimb elements (Fig S3). The shifts to lower individual forelimb element length in Apodiformes are associated with a downshift in allometric scaling relationship (Fig 2, S2). In the radius and ulna these downshifts are also associated with an increase in the slope of the allometric scaling relationship (Fig S2). The humerus displays three additional regime shifts in allometric adaptive optima: one on the branch leading to Psittacopasseres (i.e., parrots and perching birds; bayou posterior probability = 0.52), one within Ornithomimosauria on the branch leading to Ornithomimidae and Deinocheridae (bayou posterior probability = 0.63,Fig 2a) and one on the branch leading to *Ornitholestes hermanni* and Compsognathidae (bayou posterior probability = 0.34, Fig 2a). In the case of Psittacopasseres, this shift is towards a shorter humerus and this clade appears to form a distinct phenotypic optima compared to the rest of the species studied. The shift leading to Ornithomimidae and Deinocheridae is characterized by an increase in humerus length, but does not appear to form a distinct adaptive zone. These additional shifts are associated with a downshift in relative humerus length in Psittacopasseres and Compsognithidae+*Ornitholestes* and an upshift within Ornithomimosauria meaning that for a given humeri are relatively longer per a given body mass in Ornithomimidae+Deinocheiridae than compared to their ancestors.

**Fig 2:**
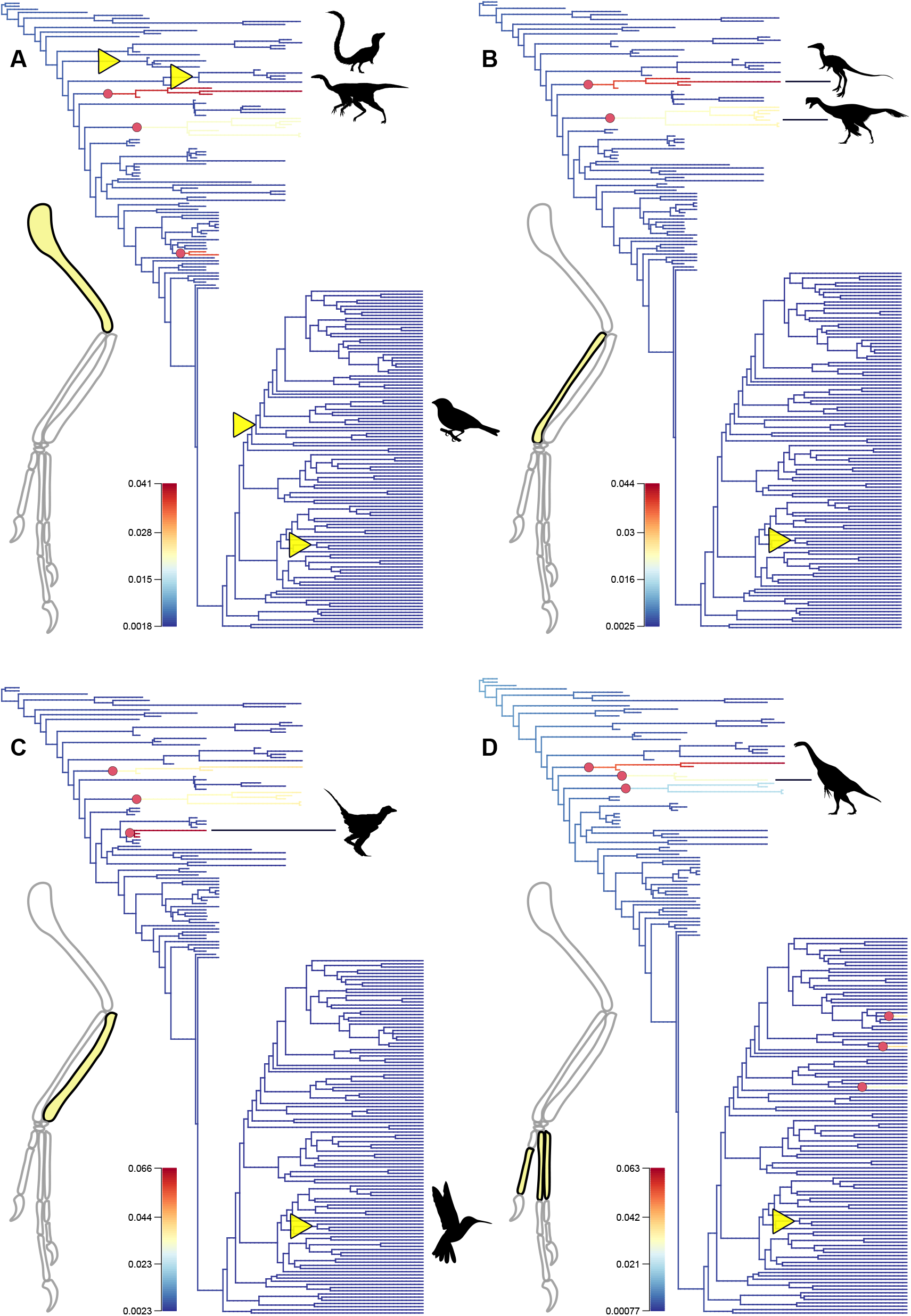
Shifts in adaptive optima (yellow triangles) and evolutionary rates (red circles) of individual forelimb bone lengths across non-avian theropods and avialans (including extant birds). A-D) Phylogenetic trees of non-avian theropods and avialans depicting the evolutionary rates for individual forelimb bone length (warmer branch colours denote faster rates of morphological evolution for the humerus (A), radius (B), ulna (C), and longest metacarpal/carpometacarpus (D). Forelimb silhouettes of *Archaeopteryx* (adapted from Middleton & Gatesy 2000) with individual bones highlighted in yellow. Sillhouettes of non-avian theropods and avialans are taken from PhyloPic.org and correspond to representative members of Compsognithidae (*Sinosauropteryx prima*, credit: Maija Karala), Ornithomimidae (*Struthiomimus altus*), Alvarezsauroidea (*Shuvuuia deserti*), Oviraptorosauria (*Oviraptor philoceratops*, credit: Ivan Iofrida), Anchiornithidae (*Aurornis xui*, credit: Gareth Monger), Apodiformes (Trochilidae), Therizinosauria (*Nothronychus mckinleyi*, credit: Scott Hartman)

There are multiple regime shifts in the adaptive optima of forelimb proportions (Fig 2). There is a regime shift in brachial index representing a shift in phenotypic optima to relatively longer proximal portion of the forelimb on the branch leading to Ceratosauria (bayou posterior probability = 0.81). There are two further regime shifts in adaptive optima in BI: one on the branch leading to Avialae (bayou posterior probability = 0.58) and one on the branch leading to Sphenisciformes (bayou posterior probability = 0.66, Fig 3). Avialans and non-avian theropods appear to occupy separate adaptive optima in BI, with avialans exhibiting lower BI scores than non-avian theropods Fig 3). Avialans display a downshift in the allometric scaling of BI relative to non-avialan theropods (Fig 3). Sphenisciformes are the exception to this rule and display much higher values for BI than any other avialan (extinct or extant), and fall well within the non-avian theropod BI adaptive peak (Fig 3). This shift is associated with an upshift in the allometric scaling of brachial index when compared to sphenisciform ancestors (Fig 3).

**Fig 3:**
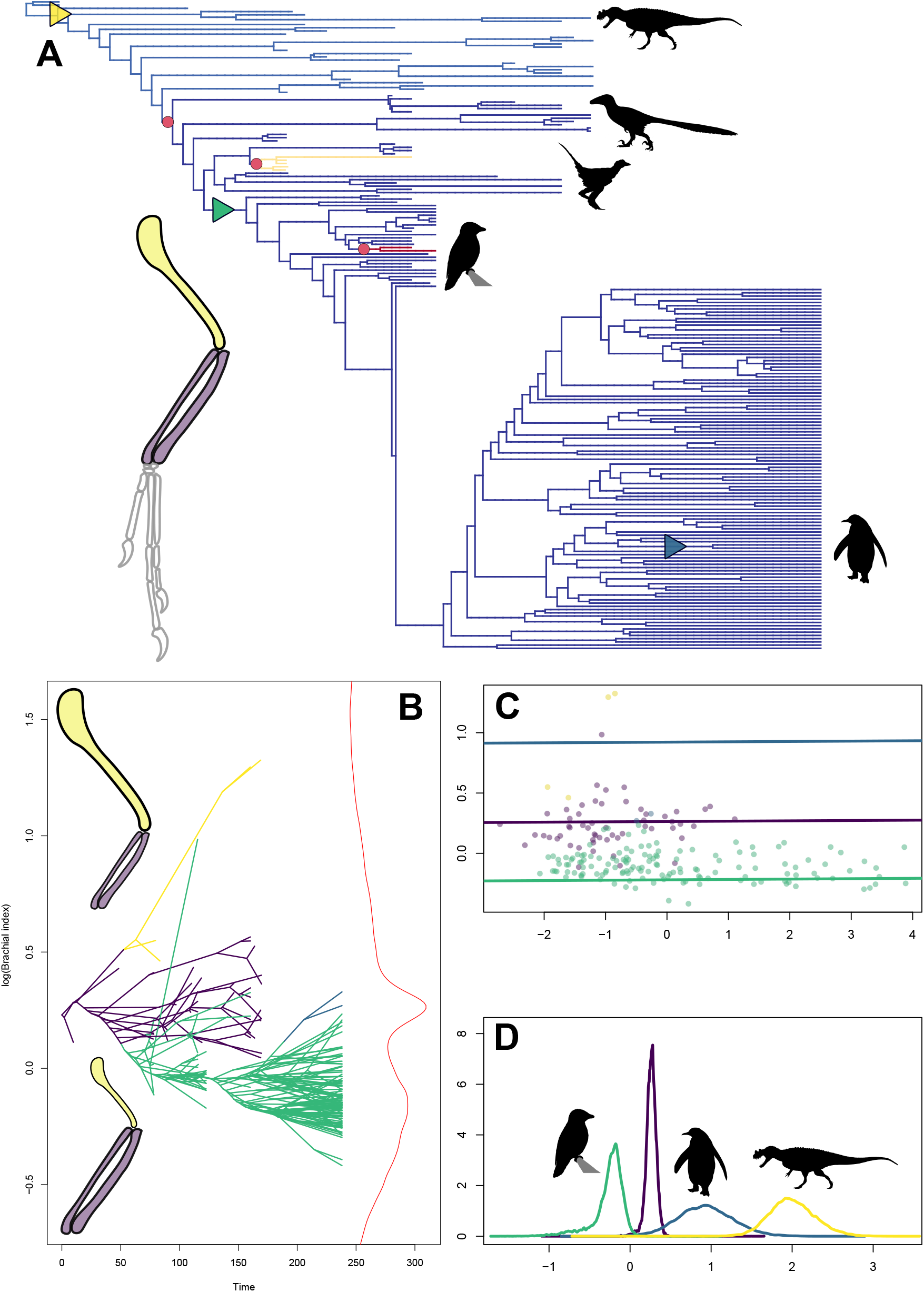
Shifts in adaptive optima (triangles) and evolutionary rates (red circles) of brachial index across non-avian theropods and avialans (including extant birds). A) Phylogenetic trees of non-avian theropods and avialans depicting the evolutionary rates of brachial index morphology (warmer branch colours denote faster rates of morphological evolution) for brachial index. Forelimb silhouette depicts the brachial index as the ratio of humerus (yellow) to radius-ulna (purple) length. B) Traitgram depicting the mode of phenotypic optimum evolution for brachial index (changes in brachial index are depicted at the extremes of the Y-axis) evolution of the last cervical vertebrae. C) Allometric scaling relationships of brachial index across non-avialan theropods and Avialae. D) Density chart for the phenotypic optima of brachial index across non-avian theropods and avialans. Triangle colours in A and line colours in B-D denote regimes of non-avian theropods and avialans where a shift in brachial index adaptive optima has occurred (yellow: Ceratosauria, purple: Neotheropoda, Green: Maniraptora (sans Alvarezsauroidea, blue: Sphenisciformes). Silhouettes are taken from PhyloPic.org and correspond to representative members of Ceratosauria (*Ceratosaurus nasicornis*, Scott Hartman), Maniraptora (*Deinonychus antirrhopus*, credit: Emily Willoughby), Anchiornithidae (*Aurornis xui*, credit Gareth Monger), Longipteryigidae (*Iberomesornis romerali*, credit: Durbed/T. Michael Keesey), Sphenisciformes (*Aptenodytes patagonicus*).

Two regime shifts in adaptive optima were observed for head length: one on the branch leading to *Mei long* (an early Cretaceous troodontid) and the other along the branch leading to *Parus atricapillus* (black capped chickadee). These shifts correspond to a decrease in BI, however neither form a new adaptive peak.

### Shifts in the evolutionary rate of neck, forelimb and head morphology

Rates of cervical morphological evolution were generally lower in avialans than in non-avian theropod dinosaurs (Fig 1). Shifts to decreased rates of centrum length evolution were observed on the branch leading to Ornithuromorpha for the middle cervical vertebrae (BAMM marginal shift probability = 0.35), and on the branch leading to Aves for the last cervical vertebrae (BAMM marginal shift probability = 0.49). The first cervical vertebrae displayed no shifts to lower rates of evolution. Shifts to higher rates of centrum length evolution are frequently observed on branches leading to clades of herbivorous non-avian theropods for the first and middle cervical vertebrae (Therizinosauria – first/middle: BAMM marginal shift probabilities = 0.93/0.83, Alvarezsauroidea – first/middle: BAMM marginal shift probabilities = 0.78/0.69, Ornithomimidae – first/middle: BAMM marginal shift probabilities = 0.55/0.44, Oviraptorosauria – first: BAMM marginal shift probability = 0.56). Additional shifts to higher rates of centrum length evolution can be observed at the base of Bohaiornithidae (first/middle: BAMM marginal shift probabilities = 0.61/0.79) and Emberizoidea (first/last: BAMM marginal shift probabilities = 0.34/0.51). None of the above shifts directly correlate to the recovered shifts in adaptive optima, however the two shifts to lower rates of centrum length evolution for the middle and last cervical vertebrae are found along the avian stem lineage, crown-ward of the avialan base. The support for the best shift configurations of centrum length evolution across all cervical vertebrae was low (<4.1% of sampled posterior distributions).

Similar to the patterns observed for centrum length rate evolution, shifts to higher rates of morphological evolution can be observed on the branches leading to herbivorous theropods across all individual forelimb elements studied (Alvarezsauroidea: BAMM marginal shift probability = 0.45-0.73, Oviraptorosauria: BAMM marginal probability = 0.35-0.93, Fig 2). Alvarezsauroidea displays the fastest rates of humerus, radius and MC2 evolution. An additional shift to higher rates of morphological evolution can be observed on the branch leading to Therizinosauria for MC2 length (BAMM marginal probability = 0.55, Fig 2). Rate shifts across all studied individual forelimb elements did not match the shifts found for their adaptive optima (Fig 2). Support for these ‘best’ configurations of rate shifts was low across all individual forelimb elements (all represented <1% of the sampled posterior distribution.

A shift to a lower rate of brachial index evolution is observed on the branch leading to Maniraptora (exclusive of Alvarezsauroidea, BAMM marginal shift probability = 0.18, Fig 3) and this shift precedes the shift in brachial index adaptive optima. Shifts to faster rates of brachial index evolution are reported along the branches leading to Anchiornithidae (BAMM marginal shift probability = 0.67) and Longipteryigidae (BAMM marginal shift probability = 0.83), Fig 3.

There appears to be little support for any rate shifts in head length across theropod and bird evolution, with the best shift configuration (accounting for 0.1% of the posterior distribution) displaying no shifts in rate. In three of the top 10 most credible rate shift configurations a shift to lower rates of head length evolution is observed on the branch leading to Avialae and these three shift configurations account for 1.23% of the posterior distribution. These rate shifts disappear after body mass is taken into account.

### Influence of allometry on shifts in neck and forelimb morphology

Ancestor-to-descendent grade comparisons indicate that the shift to a lower centrum length in Avialans that is observed across all neck vertebrae appears to be driven by decreases to centrum length outpacing changes to body mass (Tables S4, S5). The avialan ancestral grade has a mean centrum length of 3.29, 3.55 and 2.97 (log scale) and mean body mass of 1.20, 1.36 and 1.34 (log scale) for the first, middle and last cervical vertebrae respectively. The avialan descendent grade has a mean centrum length of 1.56, 1.67 and 1.52 (log scale) and mean body mass of -0.60, -0.59 and -0.58 (log scale) for the first, middle and last cervical vertebrae respectively. The difference in mean centrum length from the avialan ancestral grade to the avialan grade is -1.73, -1.89 and -1.45, whilst the difference in body mass is -1.80, -1.95 and -1.92 the for the first, middle and last cervical vertebrae respectively. The resultant 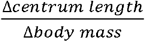 ratios across all cervical vertebrae (0.97, 0.97, 0.75) are higher (by 0.07, 0.31 and 0.01) that the upper bounds of the centrum-to-mass expectations. The 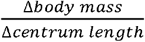 ratios across all cervical vertebrae (1.04, 1.03 and 1.33) are lower (by 0.41, 0.35 and 0.01) that the upper bounds of the mass-to-centrum expectations.

A shift to a more distally-dominated forelimb at the base of Avialae is observed for brachial index (Tables S4, S5). This shift appears to be driven by changes to body mass outpacing the changes in forelimb proportions, however there is weak support for the opposite pattern (Tables S4, S5). The avialan ancestral grade has a mean BI of 0.21 (log scale) and a mean body mass of 1.23 (log scale). The avialan descendent grade has a mean BI of -0.06 (log scale) and a mean body mass of -0.60 (log scale). The difference in BI from the avialan ancestral grade to the avialan grade is -0.28, whilst the difference in body mass is -1.84. The resultant 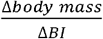 ratio for BI is higher (by 2.78) than the upper bounds of the mass-to-BI expectations. The 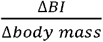 ratio for BI is only slightly higher (by 0.01) than the upper bounds of the BI-to-mass expectation.

## Discussion

We observe a concurrent shift in the adaptive optima of neck morphology (Fig 1) and forelimb proportion at the base of Avialae (Fig 3) to shorter centrum lengths in the former and to a relative increase in the distal portion of the latter. We also observe a shift in the allometric scaling of neck morphology and forelimb proportions (Figs 1, 3). Interestingly, we find no significant shifts in either adaptive optima or evolutionary rate of the length of the head (here used as a proxy for head size) across non-avian theropods or birds. These results show the neck and forelimb morphology of birds and non-avian dinosaurs display distinct evolutionary patterns as these two groups occupy separate phenotypic optima in both the neck and forelimb. Avians also exhibit slower rates of morphological evolution (Figs 1, 2) across the head, neck and forelimbs, with significant rate shifts in forelimb proportion and proximal neck morphology can occurring at the base of Maniraptora and within Ornithuromorpha respectively.

We find no support for the hypothesis that the loss of hypercarnivory in theropods released a constraint on neck evolution as no shifts in head length exist that are concurrent with (or directly proceed or precede) shifts in neck morphology at the base of Avialae. We also find very few shifts in relative head size across Theropoda more widely. Large hypercarnivorous clades are emblematic of Theropoda and the lack of significant shifts to larger relative head sizes is surprising given that hypercarniverous taxa have been shown to possess a greater selective pressure to maintain a larger relative head size (66). Phylogeny is a strong predictor theropod skull shape, more so than functional performance (67), therefore the lack of head length shifts may be a product of phylogenetic constraint. The lack of significant shifts in head size adaptive optima may also result from the measure of head size included in this study, head length. We used this metric of head size to gain access to a wider selection of theropod specimens, as relatively few are represented by a three-dimensionally preserved skull, which is necessary for volumetric reconstructions. Despite this surprising lack of adaptive optima shifts, our recovered trends in the rates of head morphological evolution match prior work and suggest that birds have a lower rate of skull evolution compared to non-avian theropods. Although head size and neck morphology may not co-evolve, recent evidence suggests that the shape of the skull (specifically the occiput) and neck morphology are integrated in extant birds due to their shared function and underlying developmental mechanisms (68), and this represents an exciting future area of research.

Our results echo the established pattern that the sustained miniaturization that occurred along the avian stem lineage heavily influenced the evolution of the Avialan appendicular skeleton (5,35,57,69) as changes to body mass outpace shifts to a more distally-dominated forelimb. Whereas forelimb evolution may be linked to selection towards a smaller body size, the shift in morphology of cervical vertebrae in basal Avialae is not, as the change in centrum length across this shift outpaces the change in body mass (Tables S2, S3). Despite not resulting in any significant shifts in head size, carnivory is still frequently associated with specific neck morphologies in both non-avian dinosaurs and extant avians (21,25,29,31) to provide stability and increased retraction forces to allow flesh to be ripped from prey items. Thus, we hypothesize that shifts in cervical morphology towards new adaptive zones is driven by the diets of basal avialans shifting away from carnivory (70,71) rather than by changes in body mass.

A shift to a more distally-dominated forelimb at the base of Avialae and to lower rates of forelimb evolution at the base of Maniraptora is supported by previous research (13,57,69). The overall proportions of the avian forelimb govern its functional capabilities. For example, an ulna that is longer than the humerus is a structural requirement of avian volant flight (72– 74). This structural requirement of powered flight has been suggested to restrict the forelimb morphology of early avialans and this restriction results in decelerated rates of forelimb morphological evolution (57). This shift in forelimb proportion is associated with a concurrent shift in vertebral morphology (centrum length), and this association has important functional consequences for the evolution of powered flight. The shift to a shorter centrum indicates an increase in cervical intervertebral stability (53,75,76), and this shift is also likely due to the structural demands imposed by the evolution of flight. The neck of extant avians plays a pivotal role in counteracting the effect of each wingbeat in order to stabilize vision during even the most complex of aerial manoeuvres (77–79). This vision stabilisation function acts as a constraint on extant avian neck morphology as soaring birds display higher levels of morphological differentiation than their continual flapping counterparts (31). Whereas the vestibulo-ocular reflex is also a critical component of avian gaze stabilisation (80), the biomechanics of the avian neck (and hence the morphology of the neck vertebrae themselves) allow for a stable vision by passively attenuating body oscillations produced from each wingbeat (78). Gaze stabilization is critical to successful locomotion in any animal (77,78,81) and is inferred here to act as a constraint on the evolution of neck morphology in the early evolution of birds.

It is this constraint of providing stable vision during flight that may be responsible for the decelerated rates of neck evolution that are observed throughout Avialae when compared to non-avian theropods. Previous work has noted that extant avian cervical morphology is generalized, with morphological adaptations only occurring to accommodate extreme neck kinematics such as those present in carnivory (31). With both early diverging avialans and extant avians showing slower rates of neck morphological evolution we suggest that the generalized morphology of the extant avian neck is a product of the constraining effect of gaze stabilization during flight. This is exemplified in the rate shift configuration of the last cervical vertebrae, whereby a shift to a much lower rate of evolution can be observed at the base of Aves (Fig. 1). The last cervical vertebrae is the most critical portion of the neck with regards to gaze stabilisation as it functions as a fulcrum for the entire cervical spine (78) thus the constraining effects of maintaining stable vision during flight should be, and are, observed in this portion of the neck. In addition, we suggest that transitions in neck morphology and forelimb proportion are not driven by shifts in the rate of morphological evolution (and vice versa) as shifts in rate and adaptive optima appear decoupled across Theropoda. The underlying drivers of evolutionary rate shifts in head size versus neck and forelimb morphology are disparate. Rate shifts in both neck and forelimb morphology are consistently found on the branches leading to herbivorous theropods (Alvarezsauroidea, Ornithomimidae, Therizinosauria and Oviraptorosauria (70)). This suggests that whereas phylogeny constrains head size rate evolution, shifts to elevated rates of neck and forelimb evolution appear adaptive and are associated with dramatic ecological shifts away from carnivory.

Heterochrony is a key mechanism of evolutionary change across vertebrates and has been found to be a vital component of dinosaur evolution (37,82–85). By analysing the changes to the intercept and slope of the allometric coefficients associated with the adaptive optima shifts of neck and forelimb morphology (Figs 1, 2, S1, S2) we can assess potential mechanisms behind the evolution of the avian neck and wing (86). Allometric shifts in adaptive optima to lower relative centrum length (Figs 1, S1) and to a lower relative brachial index are associated (Fig 3) with decreased y-intercepts and no changes to the slopes of these scaling relationships. This suggests heterochronic shifts that delay the onset of growth of these features (postdisplacement) are responsible for shifts in neck morphology and wing proportion at the base of Avialae (33,34,86). Postdisplacement is a paedomorphic process, one that results in the adult of the descendent clade (birds) resembling the juvenile stages of the ancestral clade (non-avialan theropods). Paedomorphosis is responsible for the evolution of key avian cranial characteristics (35,37) and this current work suggests that paedomorphosis is a vital mechanism behind the axial and appendicular skeleton also. Our results emphasise the importance of paedomorphosis in the emergence of avian powered flight as it provides a provides the functional adaptations (the wing) as well as the processing systems (neuroanatomical and vision stabilisation) required for life on the wing.

With concurrent shifts in adaptive optima we suggest that neck and forelimb evolution are correlated in Avialae. The grasping capability of the theropod forelimb was lost during the evolution of powered flight (72) and it has been hypothesized that the neck subsequently evolved a surrogate forelimb functionality to provide birds with a means of interacting with their environment without a grasping-capable forelimb (29–32). Some evidence for this hypothesis exists, as morphological integration between the neck and forelimb is significant in extant Aves and serves to influence the evolutionary variance of the neck (32). Neck-forelimb integration in extant avians is a keystone innovation that can allow birds to explore entirely new niches by utilizing the neck instead of the forelimb to interact successfully within these niches (32). The concurrent shifts in neck and forelimb evolution displayed here provide further support for the surrogate forelimb hypothesis and suggest that neck-forelimb evolution is correlated during the evolution of avian powered flight (29,31). Powered flight is a synapomorphy of birds and has contributed to their abundant diversity within modern ecosystems (1). This correlated evolution of the neck and forelimb to new adaptive zones represents a key innovation that has enabled birds to evolve this defining behavioral trait without losing the functionality of the forelimb.

Mosaic evolution is a core concept of avian macroevolution (6,8). This mosaicism implies that traits are organised into modules that share genetic and developmental origins. Due to these shared origins, as well as their potential shared function, traits within these modules will display a correlated response to selection. The neck and forelimb of extant birds display a coordinated evolutionary response in the form of integration, and this may have resulted from the contribution of migratory neck muscle precursor cells to forelimb muscle development (40,76,87). These developmental pathways facilitating this integration were likely present in basal avialans as they are shared across Tetrapoda (40,76) therefore the concurrent shifts in forelimb and neck evolution at the base of Avialae represent the beginning of this neck-forelimb integration. Our finding that postdisplacement is responsible for both the shifts to relative shorter centra and to a more distally-dominated forelimb provide further evidence that the neck and forelimb of avialans experience similar developmental pathways. Body mass is an important determinant of the evolution of new modules within organisms (6,15,88,89) and larger body sizes impose constraints on the evolvability of new modules within species. The miniaturization that occurred along the Avialan stem lineage may have eased these constraints and allowed for the evolution of new modules such as the one observed between the neck and forelimb. Coordinated responses of anatomical modules is a key feature of avian mosaicism and elaborations of different combinations of locomotor modules (forelimb, hindlimb and tail) have allowed birds to occupy a diverse range of niches by evolving a disparate array of locomotor behaviours (6,8,90). This study builds on recent work (31,32) to establish that avian mosaicism is not limited to locomotor modules alone (6,7). The coordinated response of neck and forelimb morphological evolution appears to be a derived feature of Avialans, initially providing them with stable vision as flight first evolved and then negating the negative consequences of losing a grasping capable forelimbs (29,31,32). In turn, this facilitated the ability for birds to interact with a multitude of different environments, and has contributed to the great taxonomic, morphological, and ecological diversity observed today.

## Acknowledgements

We would like to thank the following museum curatorial staff for their help in organizing and hosting trips to their institutions: Dr. Matthew Carrano and Amanda Millhouse at the Smithsonian Museum of Natural History, Kevin Seymour at the Royal Ontario Museum, Dr. Matt Lamanna and Amy Henrici at the Carnegie Museum of Natural History, Carrie Levitt-Bussian and Dr. Randy Irmis at the Natural History Museum of Utah, Carl Mehling, Ruth O’Leary, Suzann Goldberg and Prof. Jin Meng at the American Museum of Natural History. We would also like to thank the Leverhulme Trust (Research Project Grant award number: 182676) and the Bogue Fellowship scheme at University College London for funding this work.

## Figures

**Fig S1:**
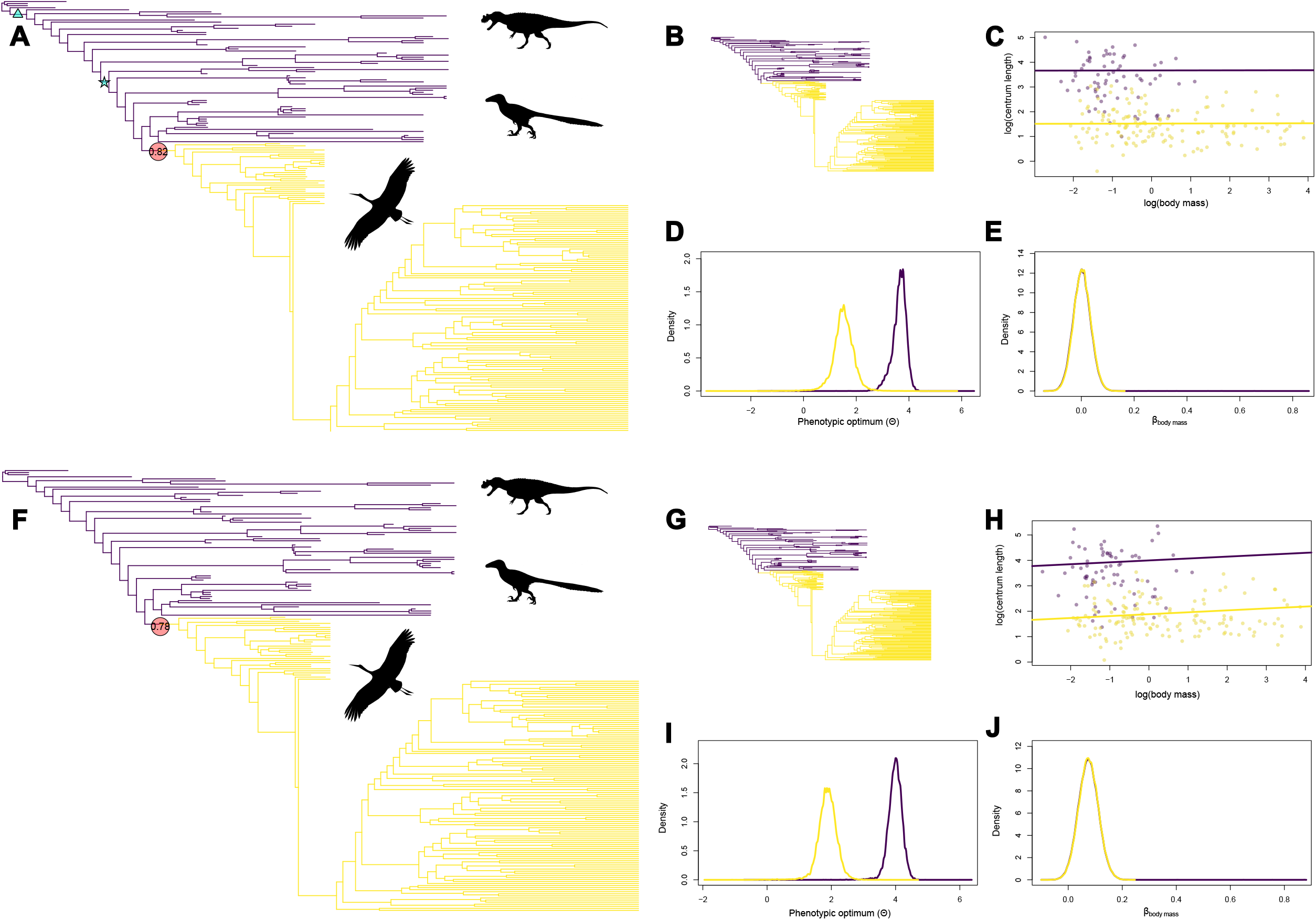
a) tree with adaptive optima shift regimes for the first cervical vertebrae, F) tree with adaptive optima shift regimes for the middle cervical vertebrae. B, G) Phylogenetic trees depicting adaptive optima regimes. C, H) Allometric scaling relationships of centrum length across the first (C) and middle (H) cervical vertebrae. D, I) Density chart displaying phenotypic optima of the first (D) and middle (I) cervical vertebrae. E, J) Density chart for assigned slope values of body mass in the associated Bayesian OU model. Across all charts, purple denotes non-avialan theropod dinosaurs and yellow denotes members of Avialae. Sillhouttes are taken from PhyloPic.org and correspond to representative members of Neotheropoda (*Ceratosaurus nasicornis*, credit: Scott Hartman), Maniraptora (*Deinonychus antirrhopus*, credit: Emily Willoughby), Aves (*Ciconia ciconia*, credit: L. Shyamal).

**Fig S2:**
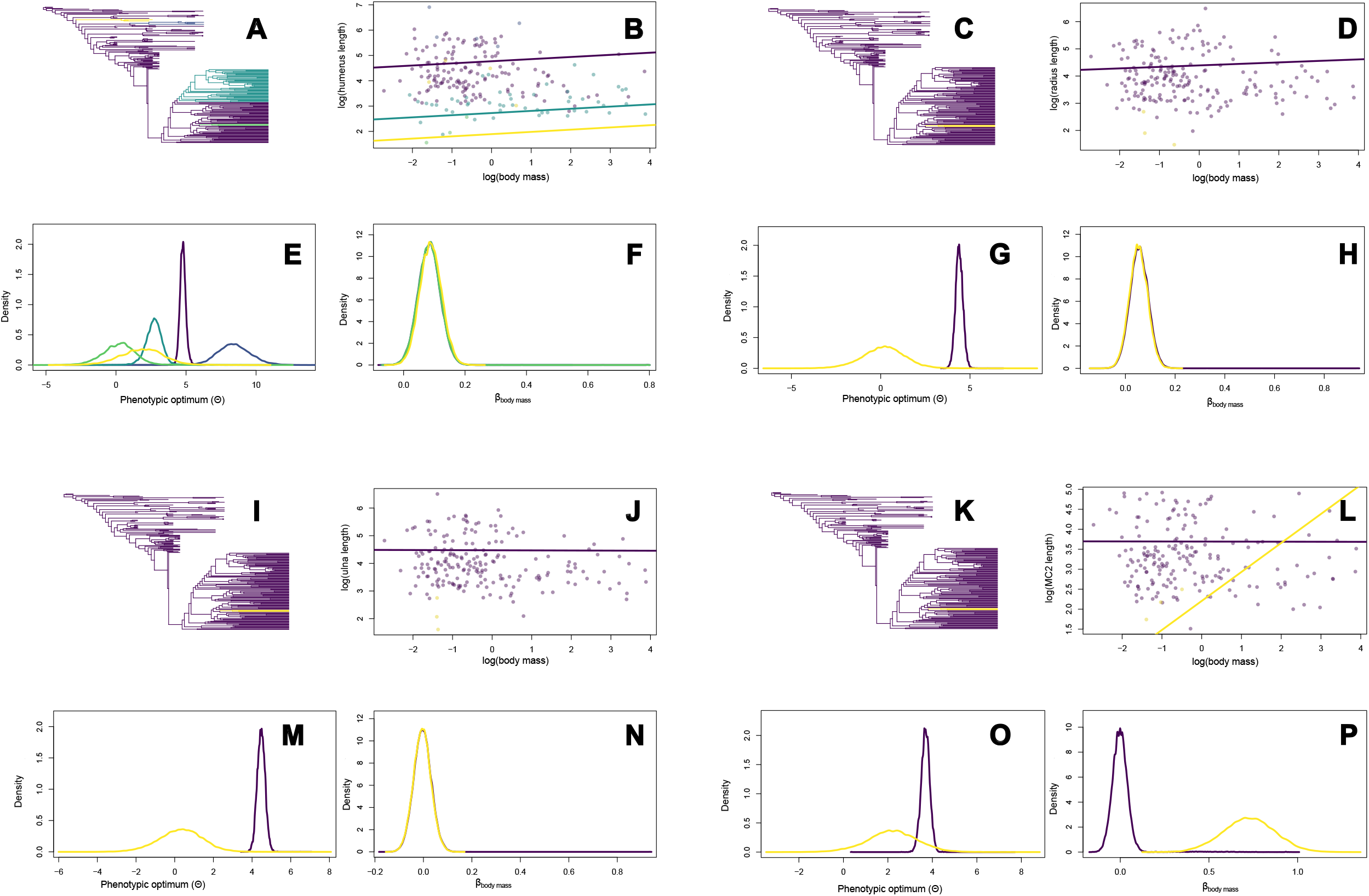
Adaptive optima regimes (A, C, I, K) and their associated allometric scaling relationships (B, D, J, L), phenotypic optima density charts (E, G, M, O) and body mass slope density charts for humerus length (A, B, E, F), radius length (C, D, G, H), ulna length (I, J, M, N) and longest metacarpal length (K, L, O, P).

