## Supplementary Information for "Correlated evolution of the neck, head and forelimb across the theropod-bird transition"

**Supplemental Figure 1**

A) tree with adaptive optima shift regimes for the first cervical vertebrae, F) tree with adaptive optima shift regimes for the middle cervical vertebrae. B, G) Phylogenetic trees depicting adaptive optima regimes. C, H) Allometric scaling relationships of centrum length across the first (C) and middle (H) cervical vertebrae. D, I) Density chart displaying phenotypic optima of the first (D) and middle (I) cervical vertebrae. E, J) Density chart for assigned slope values of body mass in the associated Bayesian OU model. Across all charts, purple denotes non-avialan theropod dinosaurs and yellow denotes members of Avialae. Sillhouttes are taken from PhyloPic.org and correspond to representative members of Neotheropoda (*Ceratosaurus nasicornis*, credit: Scott Hartman), Maniraptora (*Deinonychus antirrhopus*, credit: Emily Willoughby), Aves (*Ciconia ciconia*, credit: L. Shyamal).

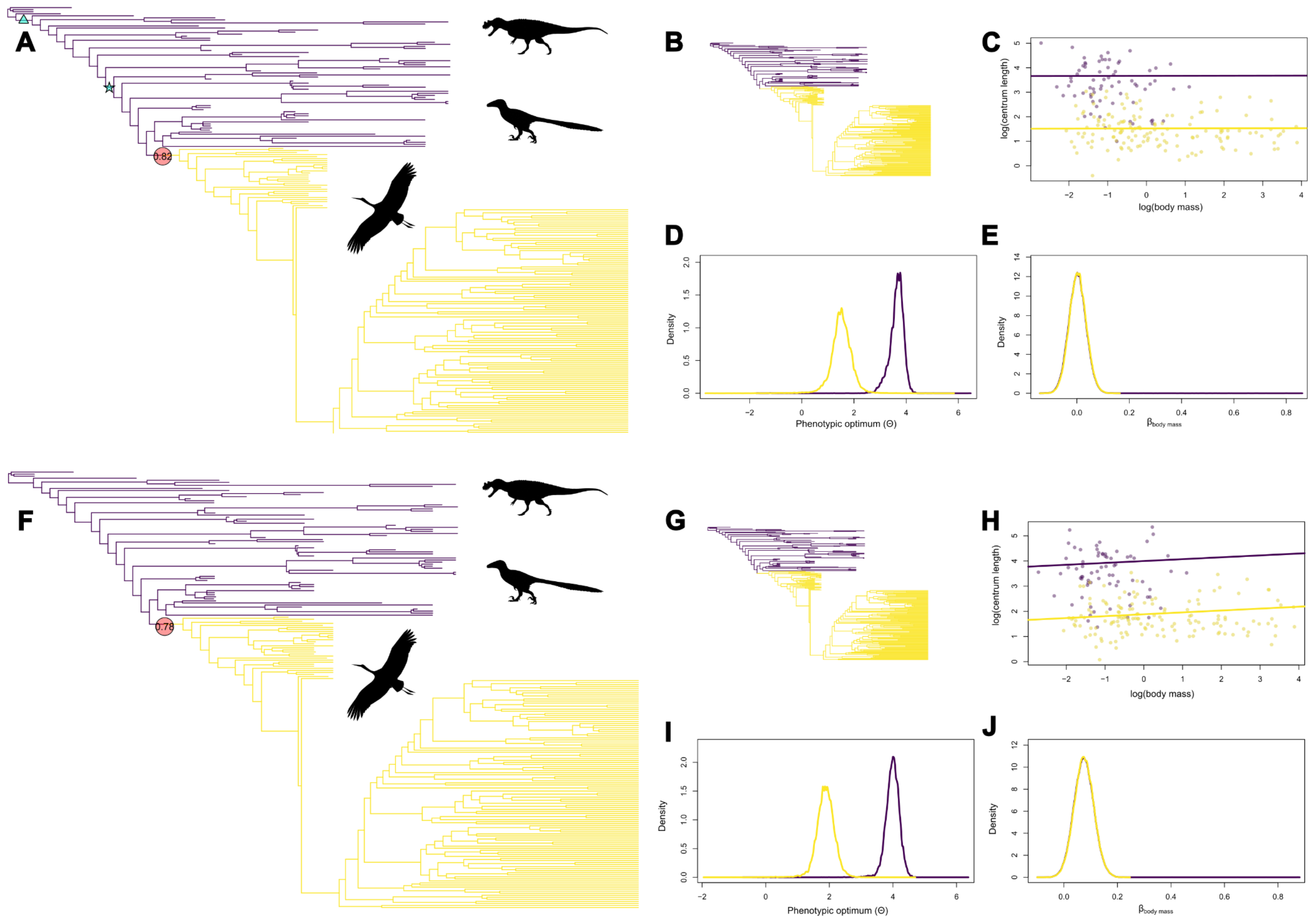
A
0.82
B
C
log(centrum length)
log(body mass)
D
Density
Phenotypic optimum (Θ)
E
Density
β body mass
F
0.78
G
H
log(centrum length)
log(body mass)
I
Density
Phenotypic optimum (Θ)
J
Density
β body mass

**Supplemental Figure 2**

Adaptive optima regimes (A, C, I, K) and their associated allometric scaling relationships (B, D, J, L), phenotypic optima density charts (E, G, M, O) and body mass slope density charts for humerus length (A, B, E, F), radius length (C, D, G, H), ulna length (I, J, M, N) and longest metacarpal length (K, L, O, P).

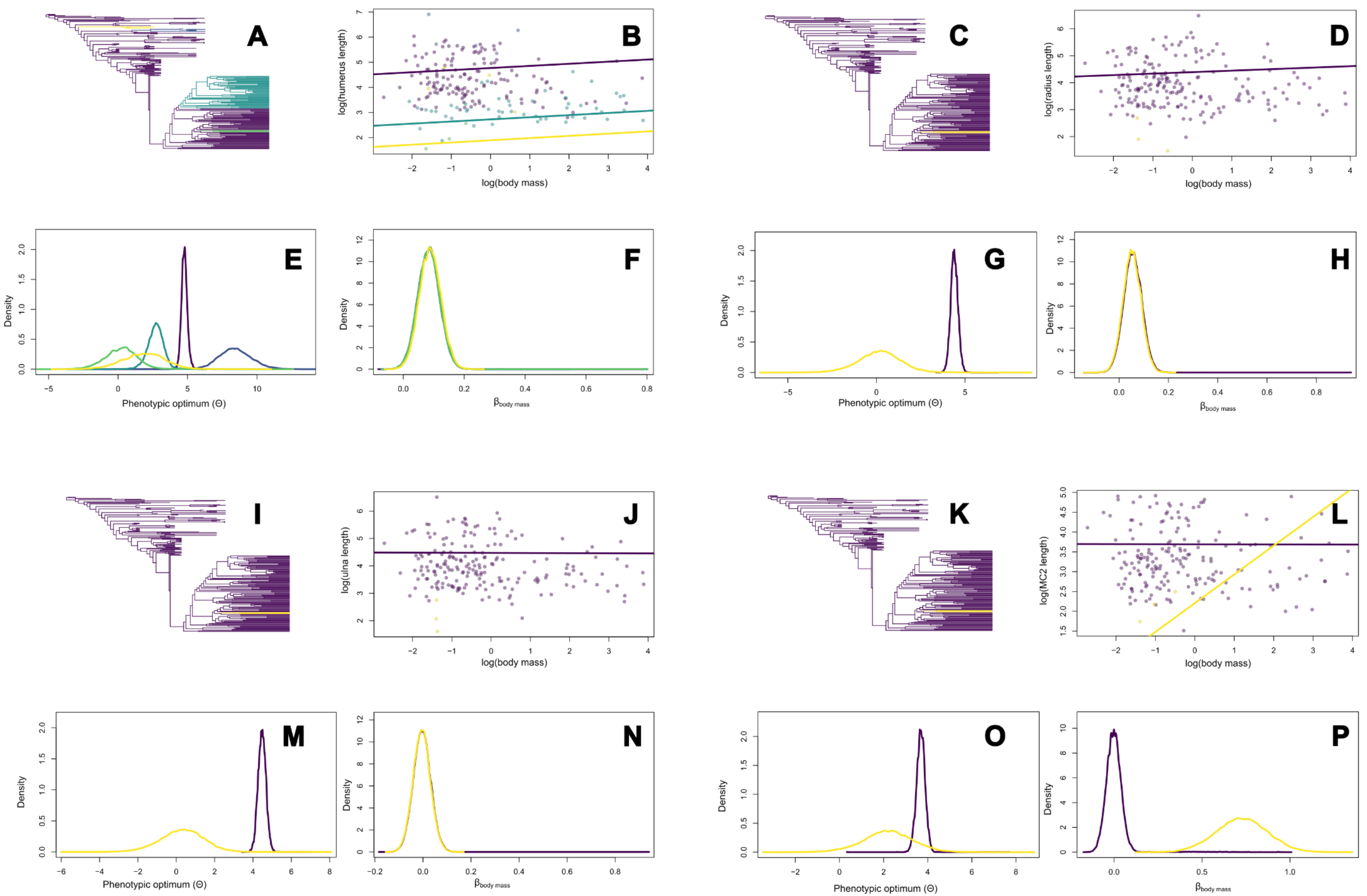
A
B
log(humerus length)
log(body mass)
C
D
log(radius length)
log(body mass)
E
Density
Phenotypic optimum (Θ)
F
Density
$\beta_{body\ mass}$
G
Density
Phenotypic optimum (Θ)
H
Density
$\beta_{body\ mass}$
I
J
log(ulna length)
log(body mass)
K
L
log(MC2 length)
log(body mass)
M
Density
Phenotypic optimum (Θ)
N
Density
$\beta_{body\ mass}$
O
Density
Phenotypic optimum (Θ)
P
Density
$\beta_{body\ mass}$

**Supplemental Table 1**

Table containing all measured data and associated metadata for each of the 224 specimens analysed as part of this study (table too large to be displayed correctly here, see attached for table).

**Supplemental Table 2**

Tables comparing Bayes Factors between three different sets of bayou models of adaptive optima shifts for each trait studied (A-H).'11' models use global intercepts and slopes for each regime across the studied trait ~ body mass relationships, 'N1' models use separate intercepts and global slopes for each regime across the studied trait ~ body mass relationships, 'NN' models use separate intercepts and separate slopes for each regime across the studied trait ~ body mass relationships (table too large to be displayed correctly here, see attached for table).

**Supplemental Table 3**

Table comparing log marginal likelihood values for a set of three models (11 = global intercepts and slopes for each regime, N1 = separate intercepts and global slopes for each regime, NN = separate intercepts and slopes for each regime) for each trait studied (table too large to be displayed correctly here, see attached for table).

**Supplemental Table 4**

Table assessing the trait-to-body mass relationship across the regimes identified in the bayou analysis by comparing phylogenetic means across ancestral and descendent regimes identified in this study

**Supplemental Table 5**

Table assessing the body mass-to-trait relationship across the regimes identified in the bayou analysis by comparing phylogenetic means across ancestral and descendent regimes identified in this study (table too large to be displayed correctly here, see attached for table).
